# Peptidoglycan remodelling improves salt resilience of *Zymomonas mobilis*

**DOI:** 10.1101/2025.10.04.680478

**Authors:** Katsuya Fuchino, Atakan Nalbant, Joe Gray, Waldemar Vollmer

## Abstract

The alpha-proteobacterium *Zymomonas mobilis* is one of the most efficient microbial producers of ethanol and has the potential to be stablished as biofuel producer at industrial scale. However, a bottleneck hindering the full use of *Z. mobilis* in biorefinery is its sensitivity to environmental stresses, including sodium chloride (NaCl), a common component present in biomass from various sources. To address this limitation, we need to deepen our understanding of the cell envelope, the crucial barrier between bacteria and external stressors. To date, the cell envelope of *Z. mobilis* has remained largely uncharacterized. Here we show that the deletion of *ctpA*, which encodes a periplasmic protease, increases the salt resilience. Salt resilience is mediated by an increased level of the peptidoglycan endopeptidase MepM, which we identified as a substrate of CtpA through comparative proteomics. Supporting this, the overexpression of MepM in the wild-type enhanced salt resilience. We also discovered that the peptidoglycan of *Z. mobilis* is O-acetylated at Mur*N*Ac residues, a modification usually associated with virulence in pathogenic bacteria. Interestingly, O-acetylation was crucial for salt resilience, supporting a role in PG growth regulation under the stress condition. Overall, this study highlights the importance of investigating cell envelope biology in *Z. mobilis* as a foundation for engineering strains with improved resilience to environmental stress and, more general, studying the cell envelopes of non-model bacteria to expand our fundamental understanding of cell function

**Importance:** Fossil fuels have negative impacts on the environment and will become limited in the next decades. Hence, alternative, sustainable energy sources need to be urgently established. Microbial fermentation of biomass for biofuel production presents a promising avenue. The Gram-negative alpha-proteobacterium *Z. mobilis* exhibits a superior capacity to convert sugars into ethanol, a clean, renewable and widely-used fuel. However, *Z. mobilis* has not been used as a first choice as a bio-fuel producer. The ethanol producer, baker’s yeast *Saccharomyces cerevisiae* serves as a model species in cell biology, but we lack fundamental understanding of the cell envelope biology of *Z. mobilis*, which would be critical to engineer strain with increased resilience. Here, we demonstrate that knowledge about cell envelope biogenesis factors in *Z. mobilis* can help engineering optimised strains that grow under conditions of bio-fuel production.

## Introduction

The Gram-negative alpha-proteobacterium *Zymomonas mobilis* has been used in the traditional production of alcoholic beverages in central America for thousands of years. It is well known for its exceptional ethanologenic physiology which enables it to convert simple sugars such as glucose into ethanol at near maximal theoretical yield and with a rapid production rate (1). Thus, *Z. mobilis* has been considered as one of the most promising microbial platforms for industrial-scale bio-ethanol production (2). Over the last two decades, metabolic engineering led to strains capable of utilising various plant feedstock for the production of ethanol and other high-value chemicals (3–5).

The central metabolism of *Z. mobilis* is well understood but little is known about the structure and biogenesis of its cell envelope, the physical barrier that separates the bacterial cytoplasm from the external environment. The Gram-negative cell envelope comprises of an outer membrane (OM), a periplasm with a thin layer of peptidoglycan (PG) and a cytoplasmic membrane. PG consists of glycan chains carrying peptides that can be cross-linked with each other, forming a mesh-like layer (6). PG surrounds the cytoplasmic membrane and, together with the firmly attached OM, provides mechanical strength to prevent bursting of the cell caused by the turgor (7). The PG also maintains the shape of the cell and the OM protect it from many toxins and antibiotics in the environment. As *Z. mobilis* remains sensitivity to various environmental stresses, which is a bottleneck for its industrial use, we need to gain a better understanding of the key cell envelope components.

One of the known stressors affecting the growth of *Z. mobilis* is common salt, sodium chloride. Even at a relatively low concentration of 225 mM, NaCl severely inhibits the growth and fermentation capacity of *Z. mobilis* for unknown reason (8, 9). Interestingly, using adaptive evolution under elevated NaCl conditions, our previous work identified a truncation of the *ctpA* gene associated with enhanced salt resilience (9). The truncation likely led to the expression of a short, inactive CtpA version lacking the whole C-terminal domain (amino acid residues 289-459). CtpA is member of the <u>C</u>arboxyl-terminal processing <u>p</u>rotease (Ctp) family of ubiquitous bacterial serine proteases. Originally, Ctp was shown to remove short C-terminal, flexible regions from its protein substrates, but further work established that it also cleaves within N-terminal domains or degrade whole substrates, regulating their cellular amount and/or activity (10). In Gram-negative bacteria, the Ctp called Prc degrades periplasmic peptidoglycan hydrolases with different cleavage sites. For example, in the model species *Escherichia coli*, Prc degrades peptidoglycan endopeptidases and lytic transglycosylases, facilitated by the Prc-activator and endopeptidase adaptor protein NlpI (11–14). If a similar regulatory mechanism exists in *Z. mobilis*, the absence of CtpA might lead to changes in the cell envelope, which could explain the observed phenotype of the salt-adapted strain with truncated *ctpA* gene.

In this study, we investigated the role of CtpA and determined the composition of *Z. mobilis* peptidoglycan. We show that modulating peptidoglycan hydrolase activity can significantly improve cell fitness under salt stress. In addition, we discovered that the peptidoglycan is O-acetylated and this modification is important for salt resilience.

## Results

### Loss of CtpA improves salt resilience in *Z. mobilis*

The previously evolved salt-resilient strain had a truncated *ctpA* gene that likely rendered the protein inactive (9). To verify that the absence of CtpA causes a salt resilient phenotype in *Z. mobilis*, we first aimed to generate a mutant strain that lacked the complete *ctpA* gene using a homologous recombination-based method (15). We readily obtained mutant colonies and confirmed the absence of the *ctpA* gene by PCR analysis. As *Z. mobilis* is a polyploid species with up to 20 chromosome copies per cell (16–19), we also checked for the deletion of *ctpA* by whole genome sequencing, which verified the deletion in all copies of chromosome (variant ratio 100%).

We next compared the growth characteristics and cell morphology of the *ctpA* deletion strain (Δ*ctpA*) and the parental strain under both salt and no-salt conditions. As previously reported, we confirmed that a low concentrations of 225 mM NaCl in the growth medium resulted in severely distorted morphology and impaired growth in the wild-type strain Z6 (8, 9); the cells displayed thick, filamentous morphology with pronounced bulging at one of the poles. In contrast, Δ*ctpA* cells grew significantly faster and to a higher optical density than the wild-type under the same salt condition (Fig. 1A). The mutant cells tended to be more elongated, but exhibited less severe polar bulging than wild-type cells (Fig. 1B). To exclude polar effects caused by the deletion of *ctpA*, we complemented the Δ*ctpA* strain by inserting *ctpA* with its upstream region into a different locus of chromosome. The complementation (strain Δ*ctpA* + *ctpA*) restored the salt sensitivity and morphology of the wild-type (Fig. 1A and 1B), verifying that the observed phenotype in the *ctpA* mutant was due to the deletion of *ctpA*.

**Fig 1.**
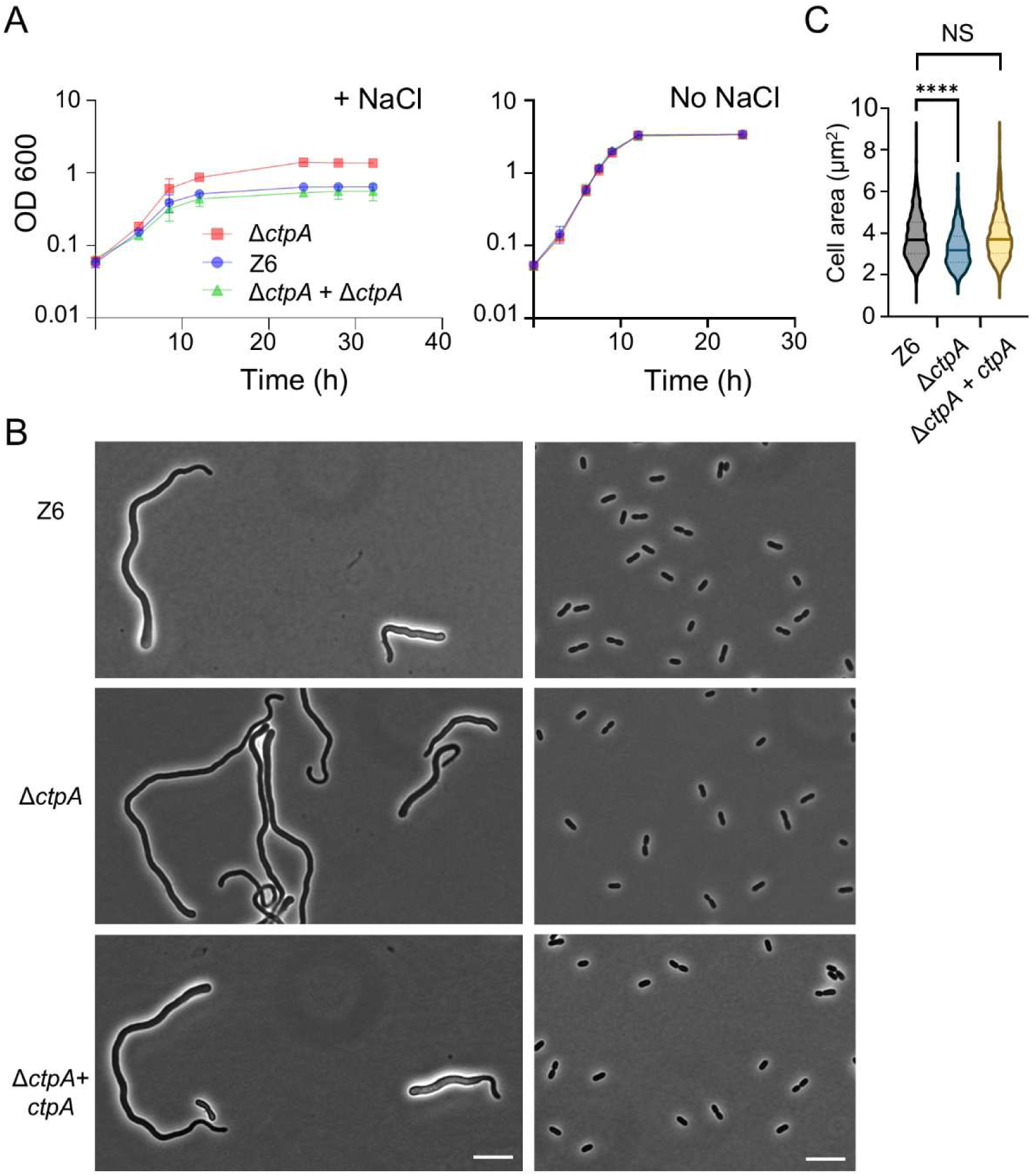
Deletion of *ctpA* improves salt resilience in *Z. mobilis*. (**A**) Growth of *Z. mobilis* strains Z6 (wild-type), Δ*ctpA*, Δ*ctpA* + *ctpA* (complementation strain) under salt stress conditions (left) and regular, no-salt conditions (right). Biological replicates N = 3. The error bars represent the standard deviation. (**B**) Phase contrast images of growing Z6, Δ*ctpA*, Δ*ctpA* + *ctpA* (complementation strain) under salt conditions (left) and regular, no-salt growth conditions (right). Scale bars, 10 μm. (**C**) Box-and-whisker plot showing the area of *Z. mobilis* cells of the three different strains grown to an OD_600_ of 0.7-1.5 under no-salt condition. The cell area was measured using MicrobeJ (39). Biological replicates N = 3. Sample size, n = 1356 cells (Z6), n = 1239 cells (Δ*ctpA*), n = 1140 cells (Δ*ctpA*+ *ctpA*). NS, not significant; ****, P ≤ 0.0001 (unpaired t-test).

Under no-salt, standard growth conditions, Δ*ctpA* and wild-type cells grew with similar rates (Fig. 1A, right side). However, a microscopic analysis revealed that the Δ*ctpA* cells were by 14.7% smaller (measured by cell area) than the wild-type cells (Fig. 1C). This phenotype was reversed by the complementation, showing that the size reduction was also due to the loss of CtpA. These findings suggest that CtpA plays a yet unidentified role in cell morphology and size, presumably by affecting the synthesis or maintenance of the cell envelope or peptidoglycan, even under no-stress conditions.

### Putative substrates of CtpA

We next asked what the underlying mechanisms of the CtpA-dependent salt sensitivity and cell size maintenance are. As CtpA is a predicted protease, we aimed to identify its substrate(s) that could influence growth and cell morphology. To this end, we performed comparative proteomics to search for putative substrates of CtpA amongst the total cellular proteins (comparing wild-type vs Δ*ctpA*, under regular or salt conditions). As expected, we did not detect CtpA in the Δ*ctpA* strain under both conditions. Substrates of CtpA are expected to be present in the periplasm and enhanced in the *ctpA* mutant (or decreased in the wild-type). Our proteomics analysis identified 42 possible substrates that were enhanced at least 1.5-fold in Δ*ctpA* cells under salt conditions (Fig. 2A, Table S3). Among these, only two proteins were also present at significantly higher amount under regular growth conditions. We identified the product of the *Z6_1677* gene as the top candidate for a CtpA substrate under both conditions (Fig. 2). *Z6_1677* encodes an M23 endopeptidase with homology to MepM in *Escherichia coli* (20.7% amino acid identity, 30.6 % similarity) (Fig. S1). Interestingly, MepM has been previously identified as a substrate of CtpA in *Pseudomonas aeruginosa* and Prc in *E. coli*, respectively (20, 21). Another protein significantly enhanced under both conditions was FtsL (encoded by *ZZ6_0468*), an essential cell division protein (Fig. 2). Other proteins enhanced only under salt conditions included three TonB-dependent proteins (encoded by *ZZ6_1483*, *ZZ6_1535* and *ZZ6_1024*), an ammonium transporter (ZZ6_0900) and a potassium transporter (ZZ6_0125) (Table S3). These proteins might contribute to the enhanced salt resistance by transporting osmolytes from the environment.

**Fig 2.**
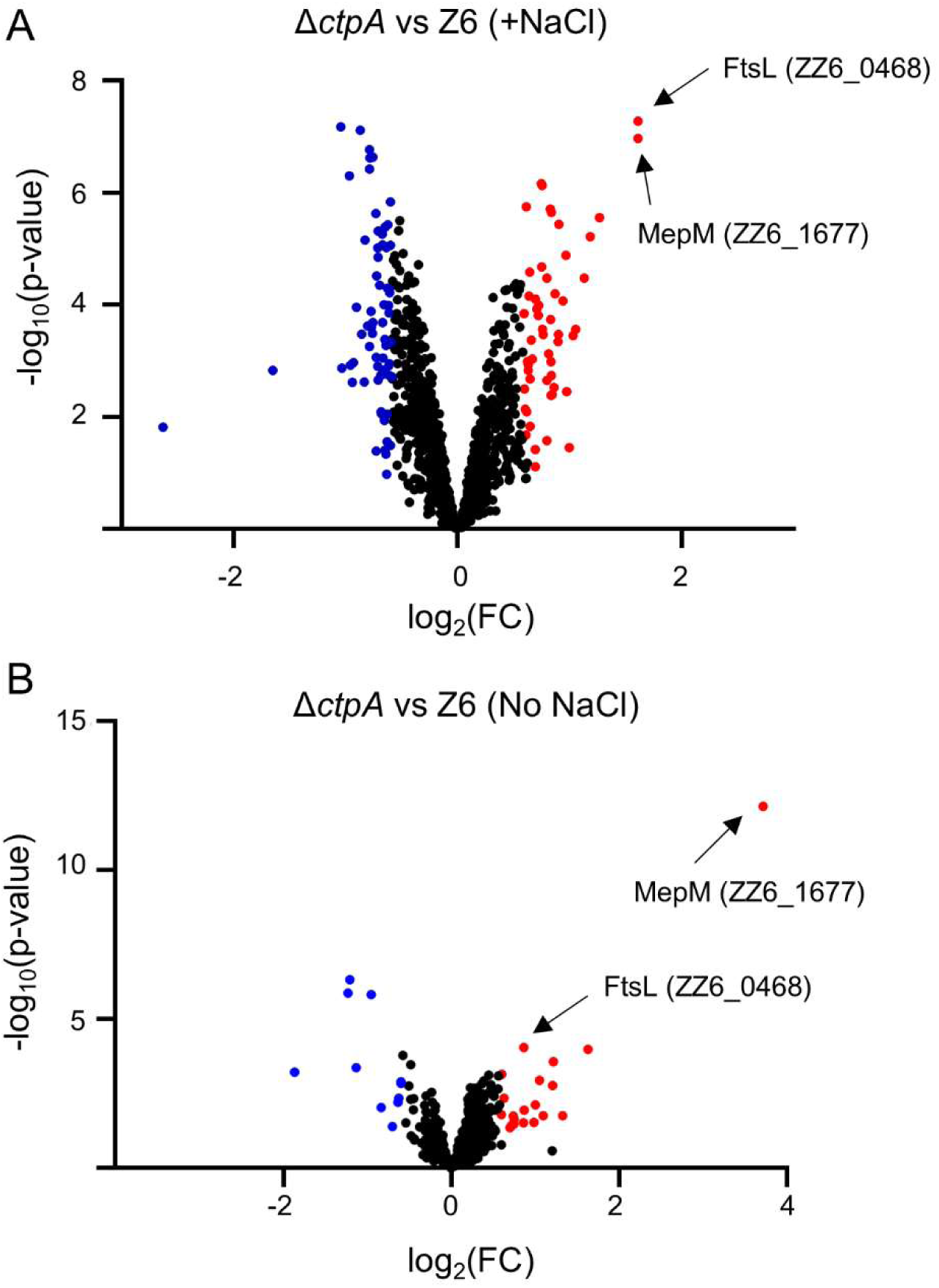
MepM and FtsL are enhanced in Δ*ctpA* under salt and no-salt conditions. Volcano plot presenting log2 fold change (FC) of abundance of individual protein (Δ*ctpA* vs Z6 under salt (**A**) and no-salt (**B**) conditions) in X-axis and its statistical significance (p-value from T-test) in Y-axis. 4 biological replicates were measured for each strain and conditions. Red indicates significantly enhanced proteins in Δ*ctpA*, blue indicates significantly reduced proteins in Δ*ctpA*. Table S3 lists enhanced proteins in Δ*ctpA* under salt conditions.

The altered cell morphology and proteomics suggested that the enhanced MepM endopeptidase level could contribute to salt resilience in Δ*ctpA*. We therefore focused on investigating the role of MepM in salt-related phenotypes.

### MepM overexpression increases salt resilience

To explore the potential role of MepM in salt resilience, we first examined the effect of *mepM* overexpression in *Z. mobilis*. We constructed a strain carrying the pBBR plasmid with the *mepM* gene under the constitutively active *pdc* promoter (Z6_1712) (22). We were unable to quantify the exact level of MepM due to the lack of specific antibodies and our inability to express a stable FLAG-tagged chromosomal version (data not shown). However, the *pdc* promoter has been used for the overexpression of other genes in *Z. mobilis* (22) hence we predict that the strain with pBBR-*mepM* has a higher MepM level than the corresponding strain with empty pBBR. Interestingly, compared to the strain with empty pBBR (Z6 pBBR), the strain carrying pBBR-*mepM* (Z6 *pdc-mepM*) exhibited enhanced growth under salt conditions (Fig. 3A, 3B), albeit the level of salt resilience did not fully reach that of Δ*ctpA* (Fig. 3A). This result strongly indicates that MepM is indeed overproduced in Z6 *pdc-mepM* and demonstrates the importance of MepM in salt resilience.

**Fig 3.**
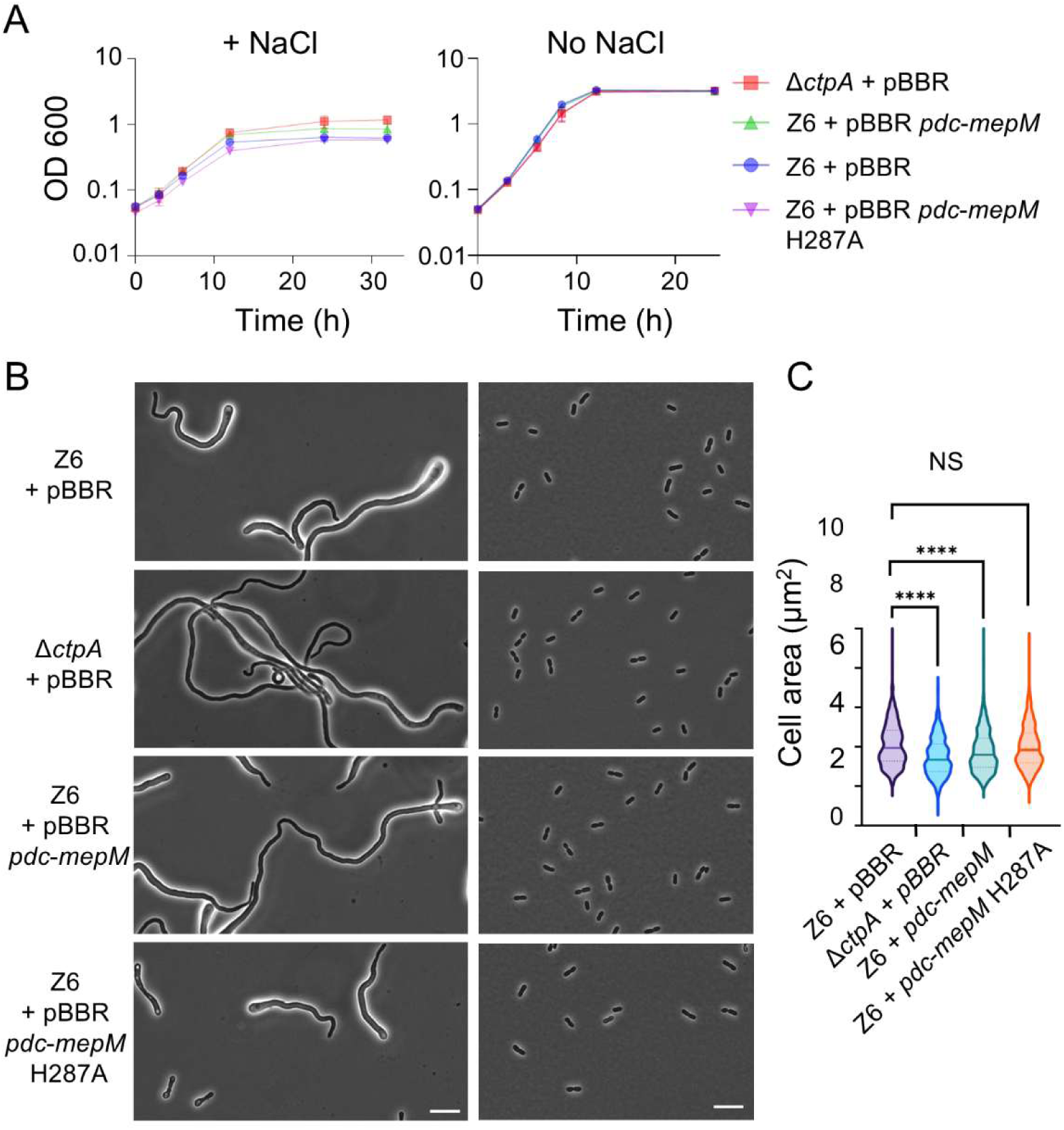
*MepM* overexpression increases salt resilience in *Z. mobilis* cells. (**A**) Growth profiles of *Z. mobilis* strains Z6 (wild-type) carrying pBBR, Δ*ctpA* carrying pBBR, Z6 carrying pBBR *pdc-mepM*, Z6 carrying pBBR *pdc-mepM* H287A under salt conditions (left) and regular, no-salt growth conditions (right). Biological replicates N = 3. The error bars represent the standard deviation. (**B**) Phase contrast images of growing, Z6 (wild-type) carrying pBBR, Δ*ctpA* carrying pBBR, Z6 carrying pBBR *pdc-mepM*, Z6 carrying pBBR *pdc-mepM* H287A under salt conditions (left) and under regular, growth conditions (right). Scale bars, 10 μm. (**C**) Box-and-whisker plot showing the area of *Z*. *mobilis* cells of the three different strains grown to an OD600 of 0.7-1.5 under no-salt condition. The cell area was measured using MicrobeJ (39). Biological replicates N = 3. Sample size, n = 585 cells (Z6 + pBBR), n = 597 cells (Δ*ctpA* + pBBR), n = 701 cells (Z6 + pBBR *pdc-mepM*), n = 593 cells (Z6 + pBBR *pdc-mepM* H287A). NS, not significant; ****, P ≤ 0.0001 (unpaired t-test).

We next investigated whether the catalytic activity of MepM is essential for the observed phenotypes in Z6 *pdc-mepM*. Previous report identified 6 critical active site residues (R286, H297, D301, H345, H376, and H378) for zinc-binding in the M23 endopeptidase ShyA, a homologue of MepM in *Vibrio cholerae* (23). Sequence alignment confirmed that 5 of these 6 critical catalytic residues (except R286 in *V. cholerae*) are conserved in *Z. mobilis* MepM. We mutated the *mepM* gene in pBBR *pdc-mepM* to express MepM (H287A) in which one of the putative critical residues, Histidine 287 is replaced by Alanine. We then transformed the plasmid into wild-type Z6 to generate strain Z6 *pdc-mepM H287A*. Under salt conditions, cells of this strain exhibited a similar poor growth phenotype as the wild-type cells, which markedly differed to the improved growth when expressing active wild-type MepM (Fig. 3A).

Under no-salt conditions, Z6 *pdc-mepM H287A* cells were similar in size as wild-type cells (Fig. 3C). This indicates that the H287A mutation abolished MepM-dependent effects and that the endopeptidase activity of MepM is essential for the salt resilience and cell size phenotypes.

To gain further functional insights into MepM in *Z. mobilis*, we aimed to delete the *mepM* gene in *Z. mobilis*. However, several attempts to delete the gene were unsuccessful, consistent with a previous systematic study using the *Z. mobilis* type-strain Z4, which identified *mepM* as an essential gene (24). To the best of our knowledge, essentiality of a single peptidoglycan endopeptidase has not been experimentally demonstrated in other bacteria. This indicates that *Z. mobilis* might possess a unique peptidoglycan hydrolase system with lower genetic redundancy, differing from that of well characterized model bacteria (25).

### Muropeptide profile of *Z. mobilis*

In *E. coli*, MepM hydrolyses DD(4–3)-crosslinks in PG peptides, contributing to making space for the incorporation of new PG during cell elongation (26). We asked if the PG composition is altered in *Z. mobilis* strains with enhanced MepM, i.e., Δ*ctpA* or Z6 *pdc-mepM*. To date the PG profile of *Z. mobilis* has not been reported and we therefore wanted to establish it for wild-type Z6, using the standard method for the Gram-negative model bacterium *E. coli* (27). Z6 cells were grown anaerobically to mid-exponential phase, harvested and boiled in the presence of 4% sodium dodecyl sulfate, followed by the recovery and purification of PG sacculi. Muropeptides (disaccharide peptide subunits) were released by the muramidase cellosyl and reduced with sodium borohydride, followed by their separation by reversed-phase high-performance liquid chromatography (HPLC) using the standard conditions for *E. coli* muropeptides (Fig. S2). This analysis showed that the major muropeptides present in the *E. coli* PG profile, disaccharide-tetrapeptide (Tetra) and bis-disaccharide-tetratetrapeptide (TetraTetra), were also present in *Z. mobilis* PG (Fig. S2). Interestingly, *Z. mobilis* PG exhibited additional major muropeptides that were absent in muramidase-digested PG from *E. coli* (Fig. S2). Mass spectrometry (MS) analysis identified these additional peaks as O-acetylated muropeptides (Fig. S2). O-acetyl groups are known to be present in the PG of several pathogenic bacteria (6). They are labile and lost at acidic and alkaline conditions, and are most stable around pH 6.0 (28). The standard HPLC buffers for *E. coli* muropeptide analysis have acidic pH (4.31 and 4.95), which likely causes the loss of some of the O-acetyl groups. We therefore repeated the PG preparation and analysis using buffers with a pH of 6.0 in all purification and analysis steps to preserve O-acetyl groups. As expected, these conditions gave rise to enhanced peaks of the O-acetylated muropeptides, but the use of HPLC buffers with a pH of 6.0 resulted in broader, less resolved peaks (Fig. S3).

The muropeptide analysis combined with MS confirmed that *Z. mobilis* possesses mature stem peptides typical for Gram-negative bacteria, L-Ala-D-iGlu-mDAP-D-Ala (Fig. 4A, S2). Crosslinked dimeric muropeptides comprised 45.9 % of the total muropeptides, which is similar as in other Gram-negative bacteria (27, 29, 30). O-acetylation was present in both monomeric and dimeric muropeptides (peak 4, 7 and 9 in Fig. 4B), affecting 55.9% of all muropeptide in the wild-type under regular growth conditions. We subsequently analysed the PG from salt-treated wild-type cells, revealing that the saline stress altered the muropeptide profile. Changes included an increase in Tri (from 1.7% to 4.1%) and Tri(OAc) (from 1.0 to 6.3 %), and a decrease in Tetra (from 19.5 to 11.9%), TetraTetra (from 5.3 to 1.8%), TetraTri(OAc) (from 3.4 to 1.8 %) and TetraTetra(OAc) (from 13.4 to 7%) (Fig. 4C). Overall, the dimers decreased from 45.9 to 43.1% and O-acetylation increased from 55.9 to 63.0% by the salt stress. Thus, salt stress induces changes in the PG in *Z. mobilis* (Fig. 4D).

**Fig. 4.**
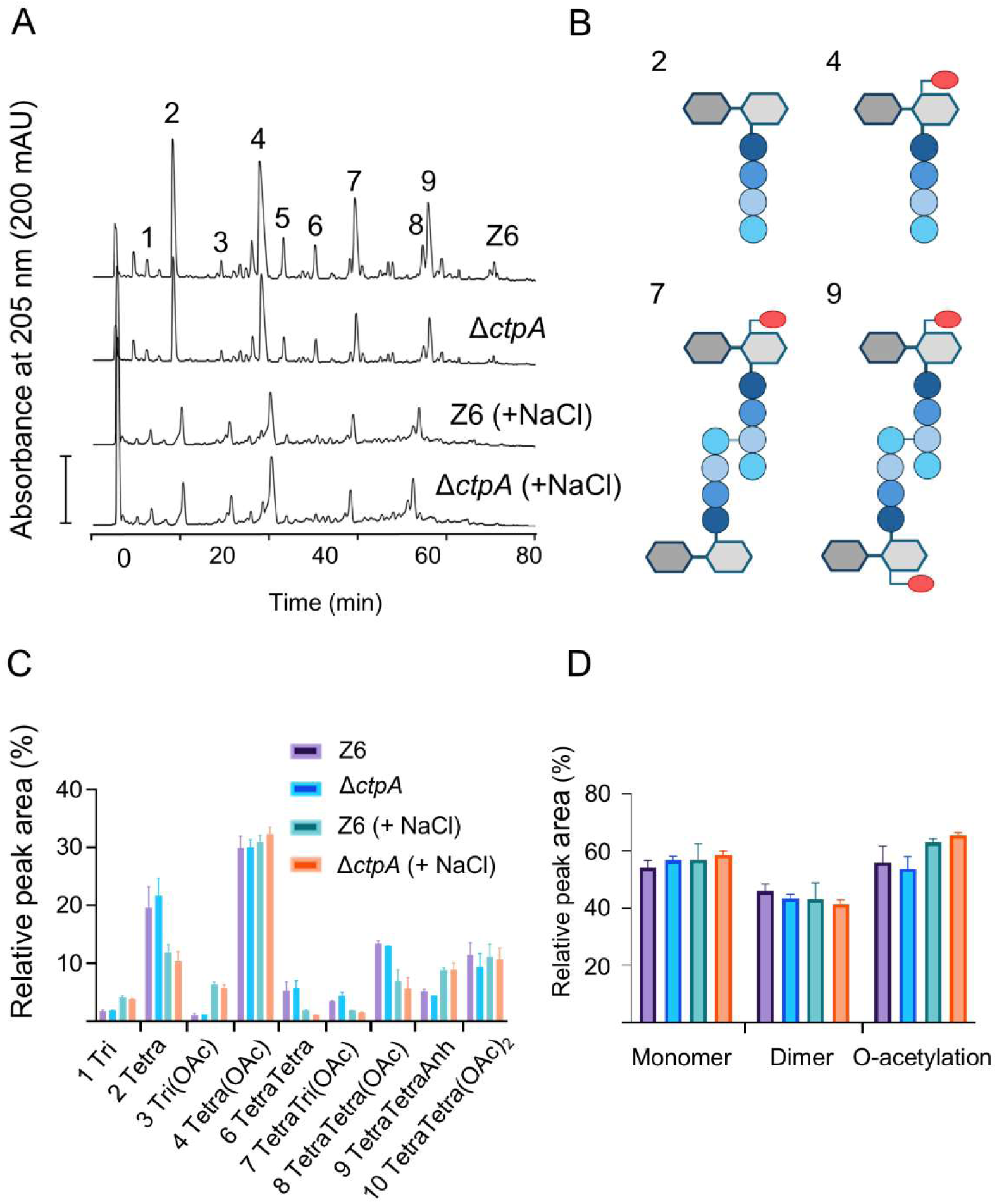
Muropeptides profile of *Z. mobilis* strain Z6 (wild-type) and Δ*ctpA*. (**A**) Cells of Z6 (wild-type) and Δ*ctpA* were grown under no-salt and salt (+ NaCl) conditions. The PG was isolated, and the muropeptides were released by cellosyl, reduced by sodium borohydride and separated by HPLC under O-acetyl group preserving conditions. Muropeptides are numbered and their structures is shown in Fig. S4. (**B**) Proposed structures of the major muropeptides released from *Z. mobilis* PG, illustrating the presence of O-acetyl groups at Mur*N*Ac in some of the muropeptides. (**C**) Relative quantification of muropeptides from Z6 and Δ*ctpA* grown under no-salt and salt conditions. Biological replicates N = 2. The error bars represent the standard deviation. (**D**) Relative proportion of monomeric, dimeric and O-acetylated muropeptides. Colours indicate the strains/growth conditions as in (C) Biological replicates N = 2. The error bars represent the standard deviation.

Next, we analysed the PG composition of the Δ*ctpA* strain for comparison. Interestingly, despite the morphological differences, the majority of muropeptides remained at similar levels between the two strains under both conditions (Fig. 4A and 4C). However, the dimers were reduced from 45.9% to 43.4% (no-salt) and 43.1% to 41.4% (salt) in Δ*ctpA* (Fig. 4D). The muropeptide profile of Z6 *pdc-mepM* was similar to that of Δ*ctpA* under regular growth conditions (Fig. S5A and S5B), with the exception of an overall 8.9% higher O-acetylation level (Fig. S5C). Under salt conditions, the dimers decreased from 42.4% to 39.0% (Fig. S5C).

### PG O-acetylation is important for salt stress response in *Z. mobilis*

Both the wild-type and Δ*ctpA* strains contained increased levels of PG O-acetylation under salt conditions, compared to no-salt conditions (Fig. 4D). In addition, Δ*ctpA* cells also contained higher PG O-Ac levels than the wild-type under salt conditions (Fig. 4D). These observations indicate that the O-Ac contents might be important for *Z. mobilis* to resist salt stress. To test the idea, we generated a mutant lacking the homologue of *patA* (Δ*patA*), which encodes a PG O-acetyl transferase present in several Gram-negative bacteria (31). The muropeptide analysis confirmed that O-acetylated muropeptides were absent in Δ*patA* cells, while the unmodified Tetra and TetraTetra were dominant in the profile (Fig. S6). As expected, the complemented Δ*patA* + *patA* strain (with *patA* on another locus on the chromosome) showed a wild type-like muropeptide profile with O-acetylated peaks (Fig. S6). This confirmed that PatA functions as a PG O-acetyltransferase in *Z. mobilis*.

We then assessed salt resilience of the *patA* mutant. Interestingly, the strain exhibited compromised growth under salt conditions while it had no growth defects under no-stress conditions (Fig. 5A). This phenotype was reversed to that of the wild-type upon complementation with the *patA* gene, excluding polar effects on adjacent genes (Fig. 5A). Interestingly, compared to wild-type, the Δ*patA* mutant cells showed a lower degree of polar bulging (Fig. 5B and 5D). Under regular, no-salt growth condition, Δ*patA* cells were slightly smaller than wild-type cells (Fig. 5C). These results indicate that O-acetylation likely affects PG synthesis and remodelling.

**Fig 5.**
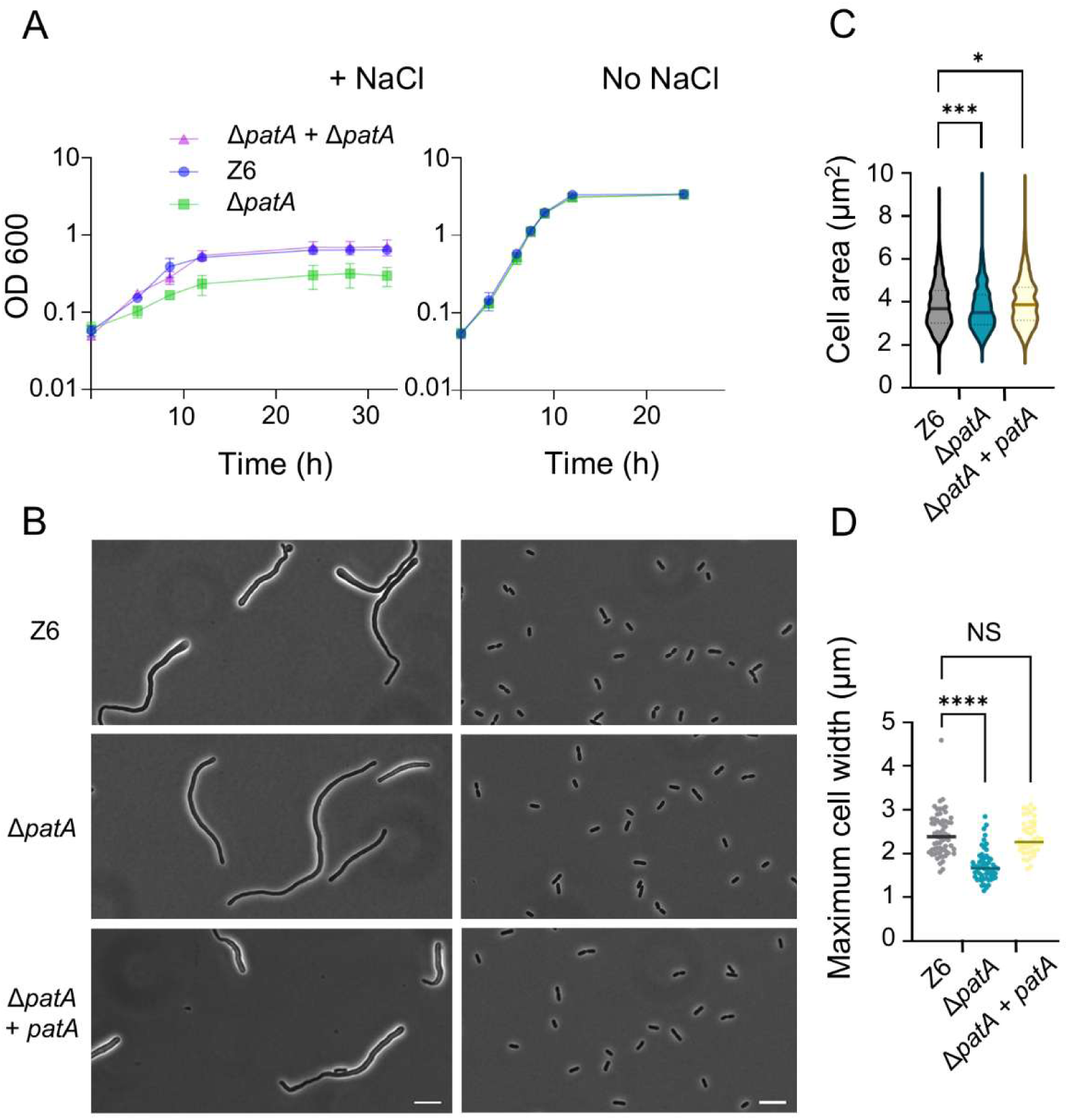
O-acetylation of PG is important for salt resilience. (**A**) Growth profiles of *Z. mobilis* strains Z6 (wild-type), Δ*patA*, Δ*patA* + *patA* (complementation strains) under salt (left) and regular, no-salt growth conditions (right). Biological replicates N = 3. The error bars represent the standard deviation. (**B**) Phase contrast images of growing, Z6 (wild-type), Δ*patA*, Δ*patA* + *patA* under salt conditions (left) and regular, no-salt growth conditions (right). Scale bars, 10 μm. (**C**) Box-and-whisker plot showing the area of *Z*. *mobilis* cells of the three different strains grown to an OD600 of 0.7-1.5 under no-salt condition. The cell area was measured using MicrobeJ (39). Biological replicates N = 3. Sample size, n = 1356 cells (Z6), n = 963 cells (Δ*patA*), n = 577 cells (Δ*patA* + *patA*). *, P ≤ 0.05; ***, P ≤ 0.001 (unpaired t-test). (D) A box-and-whisker plot presenting the maximum diameter of bulged pole of indicated *Z. mobilis* cells under salt conditions. NS, not significant, **** P ≤ 0.0001 (unpaired t-test). Biological replicates N = 3. Sample size, n = 63 cells (Z6), n = 58 cells (Δ*patA*), n = 45 cells (Δ*patA* + *patA*).

### Elevated MepM increases salt resilience independent of PG O-acetylation

Our work has so far identified two key factors contributing to salt resilience in *Z. mobilis*, elevated active MepM and O-acetylation of PG. We were curious to know if these two phenomena are linked with each other. Specifically, we wanted to examine whether MepM-dependent salt resilience depends on the presence of O-acetyl groups in the PG. To address this question, we introduced the *mepM* overexpression plasmid (pBBR *pdc-mepM*) into non-PG-O-acetylated strain Δ*patA* and assessed its growth under the salt stress condition. The analysis revealed that Δ*patA* grew better with elevated MepM level (Fig. 6A and 6B). The Δ*patA* pBBR *pdc-mepM* cells were smaller than wild-type and Δ*patA* under no-salt growth conditions (Fig. 6C).

**Fig 6.**
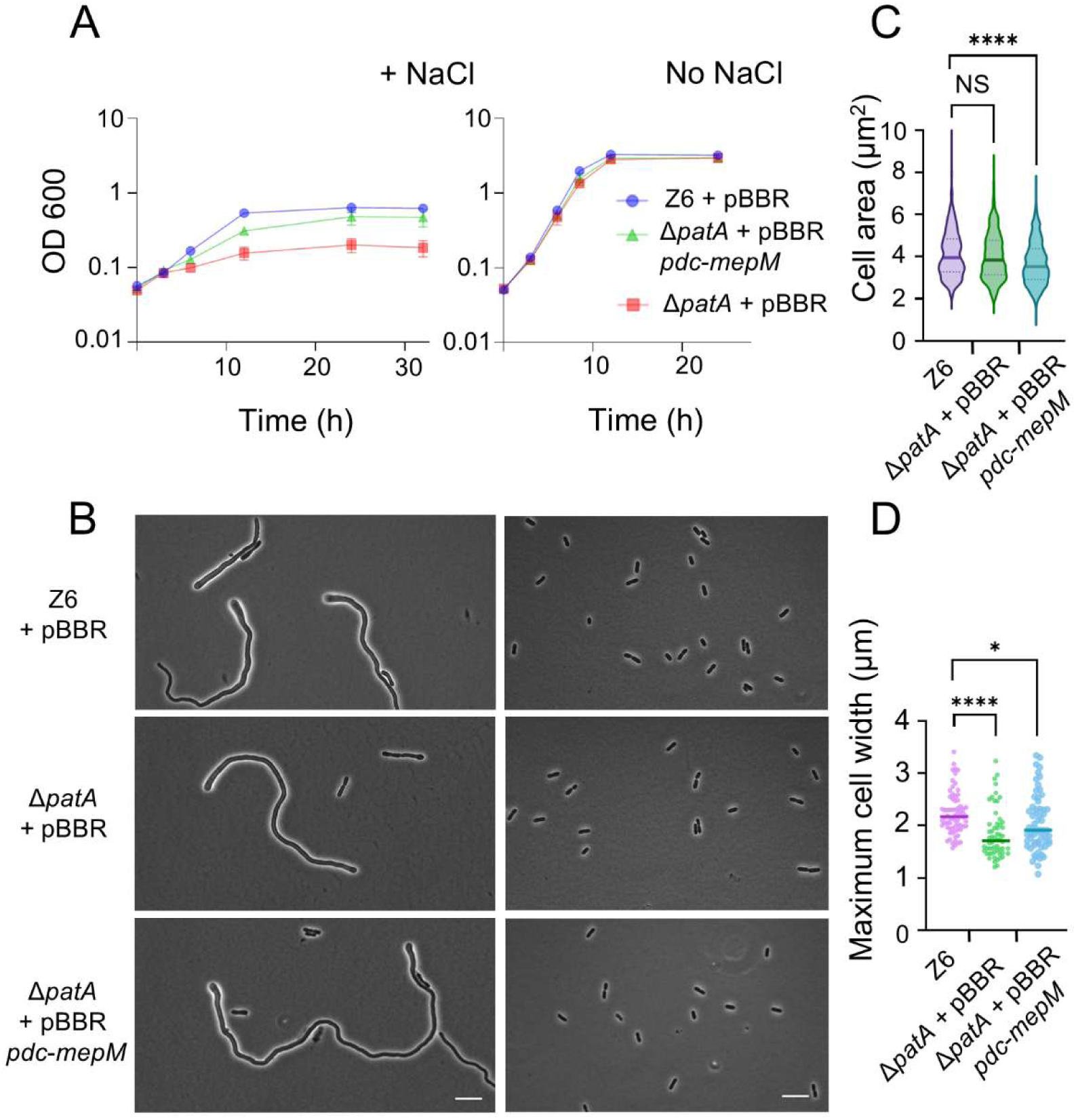
The *mepM* overexpression provides salt resilience in the *patA* mutant. (**A**) Growth profiles of *Z. mobilis* strains Z6 (wild-type) carrying pBBR plasmid, Δ*patA* carrying pBBR plasmid, Δ*patA* carrying pBBR *pdc-mepM*, under salt conditions (left) and regular, no-salt growth conditions (right). (**B**) Phase contrast images of actively growing, Z6 (wild-type) carrying pBBR plasmid, Δ*patA* carrying pBBR plasmid, Δ*patA* carrying pBBR *pdc-mepM* under salt conditions (left) and regular, no-salt growth conditions (right) Scale bars, 10 μm. (**C**) Box-and-whisker plot showing the area of *Z. mobilis* cells of the three different strains grown to an OD600 of 0.7-1.5 under no-salt condition. The cell area was measured using MicrobeJ (39). Biological replicates N = 3. Sample size, n = 585 cells (Z6 + pBBR), n = 722 cells (Δ*patA* + pBBR), n = 677 cells (Δ*patA* + pBBR *pdc-mepM*). NS, not significant; ****, P ≤ 0.0001 (unpaired t-test). (**D**) A box-and-whisker plot presenting he maximum diameter of bulged pole of indicated *Z. mobilis* cells under salt conditions. Biological replicates N = 3. Sample size, n = 70 cells (Z6 + pBBR), n = 58 cells (Δ*patA* + pBBR), n = 63 cells (Δ*patA* + pBBR *pdc-mepM*).

Furthermore, under salt condition, the ‘thin pole’-phenotype of Δ*patA* was reverted towards that of the wild-type by overexpressing *mepM* (Fig. 6D). These results indicate that MepM-mediated salt resilience and PG remodelling is independent of presence of O-Ac in PG.

## Discussion

Modulation of protein levels by the serine proteinase Ctp has been shown to be important in various biological processes across bacterial species, including virulence in pathogenic bacteria (10, 20, 32, 33), spore formation in *Bacillus subtilis* (34), and photosynthesis in cyanobacteria *Synechocystis* sp. PCC 6803 (35).

Notably, Ctp was shown to play a significant role in opposing salt stresses in two Gram-negative gamma-proteobacteria. A *P. aeruginosa ctpA* mutant grew poorer in LB *lacking* NaCl compared to the parental strain, while growth was comparable in LB with NaCl, suggesting a role of CtpA under *low-salt* conditions (20). In *E. coli*, the *prc* mutant exhibits slow growth, altered cell morphology, and frequent formation of spheroplasts under *high-salt* conditions (36), implying weakened peptidoglycan integrity in response to osmotic stress. The *prc* mutant is also sensitive to hypo-osmotic shock (37). Intriguingly, in a sharp contrast to these two species, the *Z. mobilis ctpA* mutant grows significantly better than the parental strain under high-salt conditions (Fig.1). Despite these differences, a conserved theme appears to exist in all three species: Ctp degrades the endopeptidase MepM. Our study provides first indication that the regulation of a PG endopeptidase is conserved beyond the gamma-proteobacteria and perhaps throughout proteobacteria. However, more experiments are needed to test the idea of a generalised endopeptidase degradation pathway across bacteria.

We showed that increased levels of active MepM alone significantly enhances salt resilience in *Z. mobilis* (Fig. 3), albeit level of salt resilience in Z6 + pBBR *pdc-mepM* did not fully reach that of Δ*ctpA* (Fig. 3A). This difference in salt resilience could be caused by one of the following reasons or both: 1) overproduction of MepM from the plasmid *pdc-mepM* was not at the same level as in the *ctpA* mutant, 2) other possible CtpA substrates contribute to the salt resilience phenotype. It remains unclear which of these explanations account for the observed difference.

Interestingly, the *mepM* overexpressing strains also exhibited another phenotype of Δ*ctpA*, a reduction of cell size under no-salt condition (Fig. 3C). Taken together with the proteomics data (Fig. 2), these results indicate that elevated MepM activity is a major factor underlying both the enhanced salt resilience and the reduced cell size observed in the *ΔctpA* mutant. As MepM appears to be essential for viability, it poses the intriguing question why the essential endopeptidase is constantly degraded by CtpA. One possibility is that CtpA might have a spatial control of MepM, directing its activity to specific sites within the periplasm and degrading it elsewhere.

While the exact mechanism of MepM mediated salt resilience remains unclear, it is possible that salt stress compromises the total endopeptidase activity, leading to inefficient peptidoglycan remodelling. Insufficient endopeptidase activity, possibly together with disturbed cell division, likely contributes to the elongated morphology with bulged cell shape under the stress (Fig. 1B). The elevated MepM abundance may compensate for an impaired activity of PG hydrolases needed for PG growth, explaining the enhanced growth observed in Δ*ctpA* and Z6 *pdc-mepM* cells (Fig. 7). Our study also revealed that the non-pathogenic Gram-negative *Z. mobilis* PG O-acetylates its PG and this modification is important for the salt resilience, a function never observed in previously studied bacteria with O-acetylated PG. PG O-acetylation at Mur*N*Ac has long been known to confer resistance to lysozyme in pathogenic Gram-positive bacteria (38). Gram-negative bacteria possess an outer membrane that limits the access of lysozyme to PG and O-acetylation has been proposed to modulate the activity of lytic transglycosylases (38). Under salt condition, O-acetylation might become crucial in regulating activity of lytic transglycosylases. Furthermore, we showed that salt resilience conferred by enhanced MepM does not rely on the presence of O-Ac (Fig. 6), implying that multiple factors contribute to salt resilience.

**Fig. 7.**
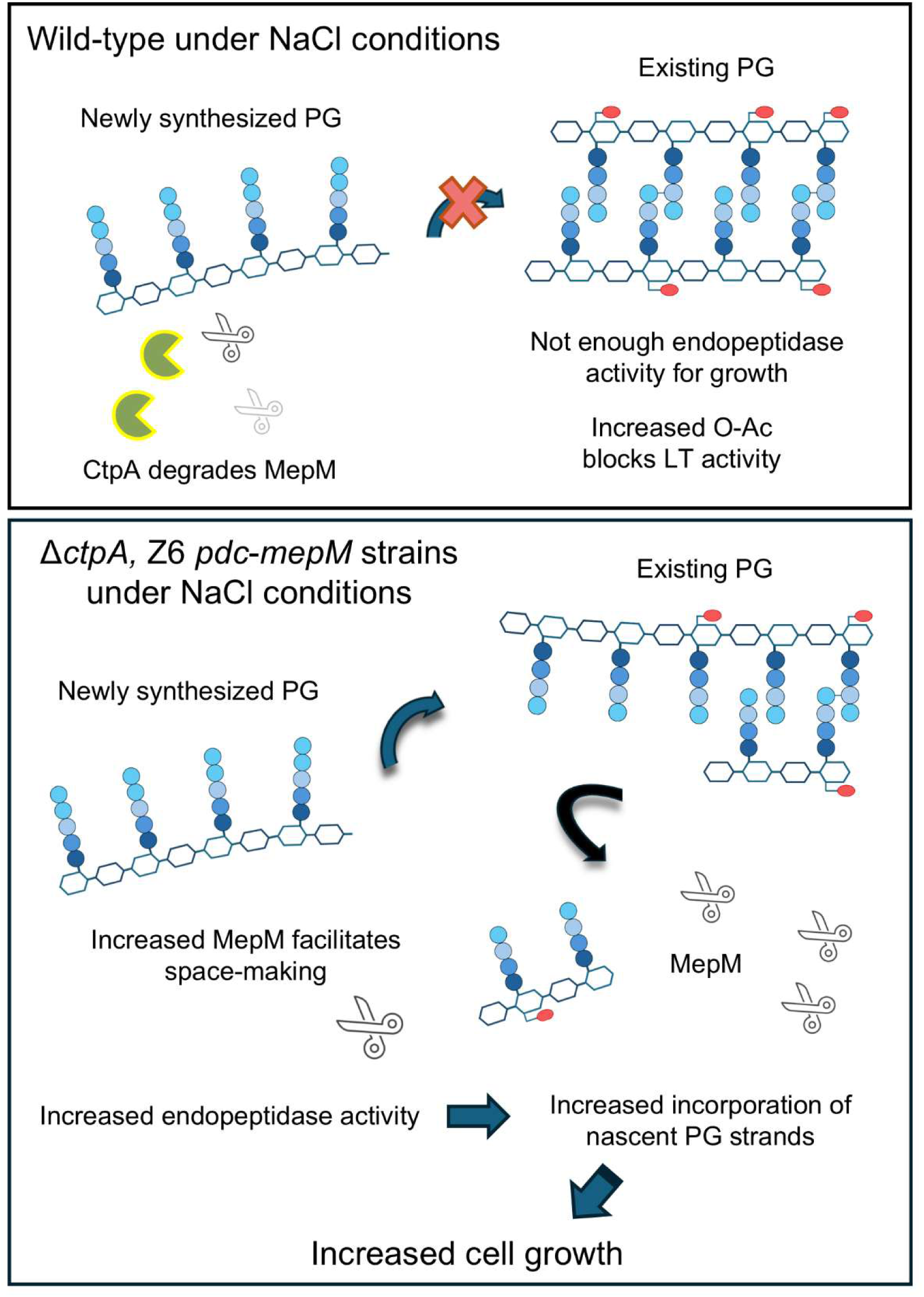
Schematic view of improved salt resilience in Δ*ctpA* and Z6 *pdc-mepM* strains. (Top) Under salt condition, wild-type cells exhibit poor growth possibly because the cells are unable to efficiently cleave crosslinked peptides in peptidoglycan. This deficiency hinders the insertion of nascent glycan chains for PG growth. The MepM level is low and in part regulated by CtpA-mediated proteolysis. (Bottom) Both Δ*ctpA* and Z6 *pdc-mepM* grow better than the wild-type under salt conditions due to higher MepM abundance and total endopeptidase activity. This allows more efficient space-making for nascent PG chains to be inserted, supporting cell growth.

Overall, this study highlights the importance of dissecting cell envelope biology as a promising strategy for engineering *Z. mobilis* strains with improved stress tolerance to enhance its potential for industrial applications. Furthermore, from a fundamental perspective, *Z. mobilis* appears to exhibit a distinct mode of cell envelope organization, which will likely offer unique insights that cannot be obtained by studying model organisms.

## MATERIALS AND METHODS

### Bacterial strains, plasmids, and growth conditions

Bacterial strains and plasmids used in this study are listed in Table S1. Oligonucleotides used in this study are listed in Table S2. *Z. mobilis* strain ATCC 29191 (Z6) was grown at 30°C in RM medium (Bacto yeast extract (5 g/L), glucose (20 g/L), NH_4_SO_4_ (1 g/L), KH_2_PO_4_ (1 g/L), MgSO_4_ (0.5 g/L) in a cap-enclosed standing tube (15 mL). For muropeptide profiling and growth assay, the RM growth medium was flashed by nitrogen gas for 45 minutes to remove oxygen.

### Construction of *Z. mobilis* strains

For genetic manipulations, we employed the method as described in (15), with a modification of the selection of cells that had undergone second homologous recombination (plasmid excision from the chromosome). To construct plasmids for gene manipulation, the upstream and downstream regions (500 – 1000 bp) of the gene of interest were amplified by Q5 DNA polymerase (New England Biolabs). The obtained cassettes were inserted into the plasmid pKK15534 by Gibson Assembly (New England Biolabs). Sequence of constructed plasmid was verified by sanger sequencing. The constructed suicide vectors were introduced into *Z. mobilis* strain Z6 via mating with *E. coli* strain WM6026. *Z. mobilis* cells carrying the integrated plasmid in their chromosomal DNA were selected by chloramphenicol 90 μg/mL in RM medium agar. The integration of the plasmid into the chromosomal DNA were confirmed by colony PCR. For excision of the plasmid, the *Z. mobilis* cells were grown in liquid RM medium without selection for overnight, and the grown cultures were subsequently selected for the loss of plasmid. In addition to fluorescence-activated cell sorting for selection, we used *Bacillus subtilis* SacB-based counter selection, a widely-used method in other bacterial system. This method was proved to be effective in *Z. mobilis* (manuscript in preparation). The genomic DNA of Δ*ctpA* and Δ*patA* mutant was sequenced by whole genome sequencing (MicrobesNG) and analysed by CLC Genomics Workbench (Qiagen).

### Microscopy and image analysis

A culture with growing *Z. mobilis* (0.5 µl) was spotted onto a 1% agarose-pad made of RM growth medium and covered by a cover-glass. Phase contrast image of *Z. mobilis* cells were captured using a Nikon Eclipse Ti microscope (Nikon) equipped with Prime sCMOS camera (Teledyne Photometrics). Image was acquired using Nikon NIS elements AR software. Image analysis was performed using MicrobeJ (39). Unpaired T-test was applied to assess statistical significance.

### Proteomics

Samples for proteomics were prepared as follows. *Z. mobilis* strains were grown anaerobically in RM medium cap-enclosed standing tubes (15 mL) to an OD_₆₀₀_ of 1.0. For salt stress conditions, RM medium was supplemented with 225 mM NaCl.

Cells were harvested by centrifugation at 15,000 ×g for 1 minute at 4°C, washed once with ice-cold phosphate-buffered saline (PBS), and cell pellets were stored at −80°C until proteomics analysis. Frozen cell pellets were lysed in cell lysis buffer containing 5% sodium dodecyl sulfate (SDS, Sigma-Aldrich), 50 mM triethylammonium bicarbonate (TEAB, Sigma-Aldrich), pH 8.5. Samples were sonicated on ice using a probe sonicator (3 × 30 seconds pulses with 30 seconds intervals) to ensure complete protein extraction and kept on ice between processing steps. Protein concentration was determined using the Pierce BCA Protein Assay Kit (Thermo Fisher Scientific) with bovine serum albumin as a standard, performed in triplicate and averaged. Protein amounts were normalized to 50 μg per sample, and sample volumes were adjusted to 23 μL with lysis buffer when necessary for consistent downstream processing.

Proteins were reduced with tris(2-carboxyethyl)phosphine (TCEP, Sigma-Aldrich) at a final concentration of 5 mM and incubated at 55°C for 15 minutes, then cooled to room temperature. Alkylation was performed with Iodoacetamide (IAA, Sigma-Aldrich) at a final concentration of 20 mM and incubated at room temperature for 15 minutes. Following reduction and alkylation, samples were acidified to a final concentration of 2.5% phosphoric acid (85% wt. in H_2_O, Sigma-Aldrich). Acidified samples were loaded onto S-Trap micro spin columns (Protifi) and digested according to the manufacturer’s protocol using sequencing-grade modified trypsin (Promega Gold) at a 1:10 enzyme-to-protein ratio (w/w) and incubated for 2 hours at 47°C. After digestion, samples were loaded unto EVOTIPS following the EVOSEP tip loading protocol. Liquid chromatography-tandem mass spectrometry (LC-MS/MS) analysis was performed using an EVOSEP ONE liquid chromatograph (EVOSEP) coupled to a timesTOF HT mass spectrometer (Bruker Daltonics) via the CaptiveSpray source. Chromatographic separation was achieved using an IonOpticks Aurora Elite 15 × 75 mm C18 UHPLC column operated in Whisper Zoom configuration. The EVOSEP 20SPD method was employed with mobile phase A consisting of aqueous (Sigma-Aldrich, UHPLC Grade) 0.1% formic acid and mobile phase B consisting of 100% acetonitrile (Sigma-Aldrich, UHPLC Grade) with 0.1% formic acid. The gradient profile ramped mobile phase B from 2% to 40% over 60 minutes at a flow rate of 200 nL/min during the gradient phase. Mass spectrometric analysis was performed in positive ion mode using data-independent acquisition (DIA) with a method optimized using the pydiAID algorithm (40). The ion source was operated with the following parameters: capillary voltage, 1.6 kV; dry gas flow, 3 L/min; dry gas temperature, 180°C. DIA acquisition was performed with precursor ion isolation windows spanning m/z 400-1000 with variable window widths optimized for even precursor distribution. The accumulation and ramp time were set to 100 ms per scan, resulting in a total cycle time of approximately 1.7 s. Trapped ion mobility spectrometry (TIMS) was enabled with 1/K₀ values ranging from 0.6 to 1.6 V·s/cm². The diaPASEF method employed for the LC-MS analysis of the samples is provided alongside the raw files in the PRIDE repository. Raw mass spectrometry data were processed using DIA-NN version 2.0.1 (41) with a predicted spectral library generated from the *Z. mobilis* ZZ6 database and contaminant database (cRAP). The following parameters were applied: precursor and protein false discovery rate (FDR) set to 0.01, maximum number of variable modifications set to 1, maximum missed cleavages set to 1, mass accuracy tolerance set to 15 ppm for both MS1 and MS2, N-terminal methionine excision enabled, and tryptic digest specificity (K*, R*).

Variable modifications included oxidation of methionine (UniMod:35, +15.994915 Da) and N-terminal acetylation (UniMod:1, +42.010565 Da). Cysteine carbamidomethylation was applied as a fixed modification. Match-between-runs (MBR) was enabled for enhanced peptide identification across samples. Protein grouping was performed using implicit protein grouping based on protein names, and contaminant proteins tagged with “cRAP-” were excluded from normalization and quantification. Peptidoform scoring and RT profiling were enabled to improve identification confidence. Results of the proteomics, as well as the raw data and LC-MS methods (embedded in the raw data files) are made available on ProteomeXchange repository (42) via PRIDE database (43). Prediction of protein localisation was assessed using Signal IP – 6.0 (44)

### Muropeptide analysis

A *Z. mobilis* culture (400 mL) was anaerobically grown to an optical density of 0.6 – 0.8. The cells were harvested by centrifugation, resuspended in ice-cold PBS and subsequently boiled in 4% SDS. Purification of peptidoglycan and high-performance liquid chromatography (HPLC) was performed as described (27, 45), with a modification of use of buffers and mobile phase with a pH of 6.0 to preserve O-acetyl groups. HPLC buffer B contained 15% methanol instead of 30% methanol as in (45). Mass spectrometry (MS) analysis for identification of muropeptide was performed as described in (45).

## Acknowledgements

We thank Prof. Patricia J. Kiley (University of Wisconsin–Madison) for donating the plasmid pKK15534. We thank Christodoulos Astraios for constructing the strain *patA* + *patA* and Jacob Biboy for assistance with the HPLC analysis. This work was supported by Newton International Fellowship (The Royal Society) to K.F. (NIF\R1\221190)

## Notes

### Competing Interest Statement

The authors have declared no competing interest.

