## Supplementary material for "Peptidoglycan remodelling improves salt resilience of *Zymomonas mobilis*": Supp Fig. S1-S6 and Supp table S1-S3

### Supplemental figures

```
MepM_Ecoli_K12         MQQIARSVALAFNNLPRPHRVMLGSLTVLTLAVAVWRPYVYHRDATPIVKTIELEQNEIR  60

MepM_Z.mobilis_Z6      ------------------------------------------------------------  0

MepM_Ecoli_K12         SLLPEASEPIDQAAQEDEAIPQDELDDKIAGEAGVHEYVVSTGDTLSSILNQYGIDMGDI  120

MepM_Z.mobilis_Z6      ------------------------------------------------------------  0

MepM_Ecoli_K12         TQLAAADKELRNLKIGQQLSWTLTADGELQRLTWEVSRRETRTYDRTAANGFKMTS---E  177

MepM_Z.mobilis_Z6      -----------------------SHAAT-------LHQKKSAAFH-TSSSHFSSSSKTAS  29

                                              :  .        : :::: ::. *::. *. :*   .

MepM_Ecoli_K12         MQQGEWVNNLLK-GTVGGSFVASARNAGLTSAEVSAVIKAMQWQMDFRKLKKGDE-FAVL  235

MepM_Z.mobilis_Z6      SKTGHFFLAQHQGGRYGSHY-AMSRNA-------------SRYAVAHRSFYHGFQPISID  75

                        : *.:.    : *  *. : * :***              :: : .*.: :* : :::

MepM_Ecoli_K12         MSREMLDGKREQSQLLGVRLRSEGKDYYAIRAEDGKFYDRNGTGLAK----GFLRFPTAK  291

MepM_Z.mobilis_Z6      PSPQMVNASQEEPI-------NRDNQYHKLFVS----WAKADTAQTEKAAVVPSATPVNS  124

                        * :*::..:*:         ...::*: : ..    : : .*. ::         *. .

MepM_Ecoli_K12         QFRISSNFNPRRTNPVTGRVAPHRGVDFAMPQGTPVLSVGDGEVVVAKRSGAAGYYVAIR  351

MepM_Z.mobilis_Z6      RFLLTSPFG-VRSDPFHRHAAMHAGVDLAAPYGTPVYASADGVVDRAGGASGYGNLVEID  183

                       :* ::* *.  *::*.  :.* * ***:* * **** : .** *  *  :.. *  * *

MepM_Ecoli_K12         HGRSYTTRYMHLRKILVKPGQKVKRGDRIALSGNTGRSTGPHLHYEVWINQQAVNPLTAK  411

MepM_Z.mobilis_Z6      HGHMIQTRYGHLSRILVHEGQAIKRGDLIALMGSTGRSTGSHLHYEVRIQGEAVNPVPFL  243

                       **:   *** ** :***: ** :**** *** *.****** ****** *: :****:

MepM_Ecoli_K12         LPRTEGLTGSDRREFLAQ--AKEIVPQLRFD---       440

MepM_Z.mobilis_Z6      AVDSYHMAMLDNVSAGAQGGPEKDVKDVKVHHHK       277

                          :  ::  *. .  **   :: * :::..
```

Fig. S1 Sequence alignment of *Z. mobilis* MepM with the sequences of *E. coli* K12 MepM, analysed by Pairwise Sequence Alignment (EMBOSS needle)

A

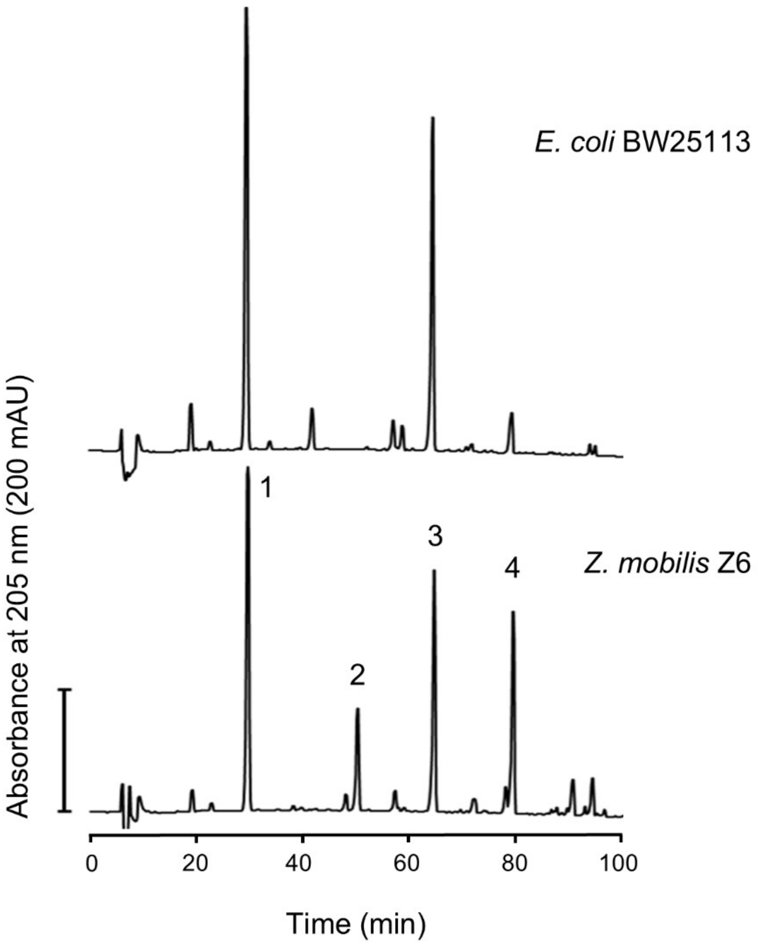


B

| Peak No. | Muropeptide | Neutral mass (amu) of the reduced form | |
|---|---|---|---|
| | | Observed | Theoretical |
| 1 | Tetra | 941.10 | 941.41 |
| 2 | Tetra(OAc) | 983.17 | 983.42 |
| 3 | TetraTetra | 1864.42 | 1864.80 |
| 4 | TetraTetra(OAc) | 1907.62 | 1906.82 |

Fig. S2 Comparison of *E. coli* and *Z. mobilis* muropeptide profiles

(A) HPLC chromatogram of reduced muropeptides from *E. coli* BW25113 (top) cells and *Z. mobilis* Z6 cells (below). The used HPLC method is as described in (27). The labelled peaks are presented in the panel B. (B) Reduced muropeptides from *Z. mobilis* cells identified by Mass spectrometry. Theoretical masses were calculated using ChemDraw™ (PerkinElmerInformatics,UK)

A

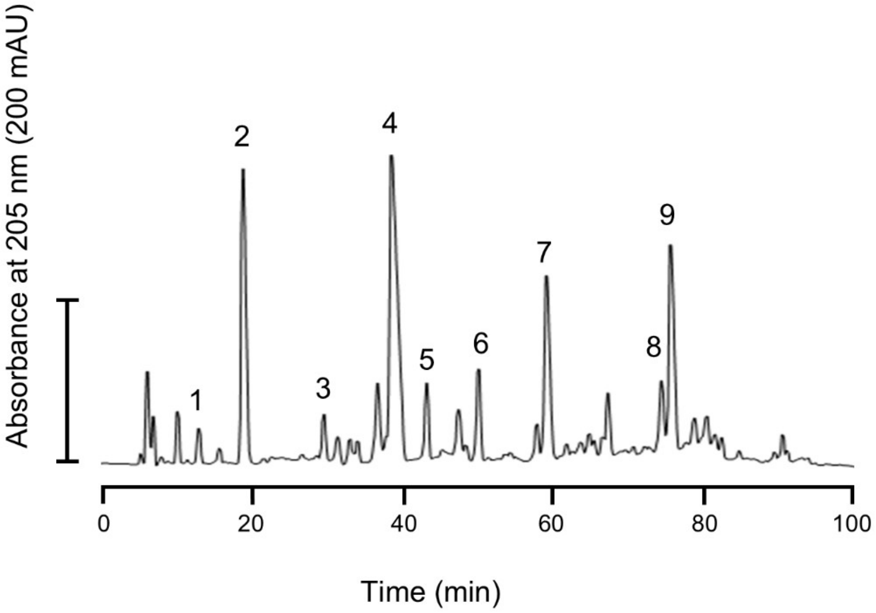


B

| Peak No. | Muropeptide | Neutral mass (amu) of the reduced form | |
|---|---|---|---|
| | | Observed | Theoretical |
| 1 | Tri | 870.06 | 870.37 |
| 2 | Tetra | 941.11 | 941.41 |
| 3 | Tri(OAc) | 912.14 | 912.39 |
| 4 | Tetra(OAc) | 983.25 | 983.42 |
| 5 | TetraTetra | 1864.66 | 1864.80 |
| 6 | TetraTri(OAc) | 1835.92 | 1835.78 |
| 7 | TetraTetra(OAc) | 1907.14 | 1906.82 |
| 8 | TetraTetraAnh | 1845.80 | 1844.78 |
| 9 | TetraTetra$(OAc)_2$ | 1949.20 | 1948.83 |

Fig. S3. Identification of *Z. mobilis* muropeptides

(A) HPLC chromatogram of reduced muropeptides from *Z. mobilis* Z6 cells. HPLC method was modified from (27) to preserve O-acetylated muropeptides. The labelled peaks are presented in the panel B. (B) Reduced muropeptides obtained from *Z. mobilis* cells. Theoretical masses were calculated using ChemDraw™ (PerkinElmerInformatics,UK). Muropeptides are numbered and their structures shown in Fig. S3

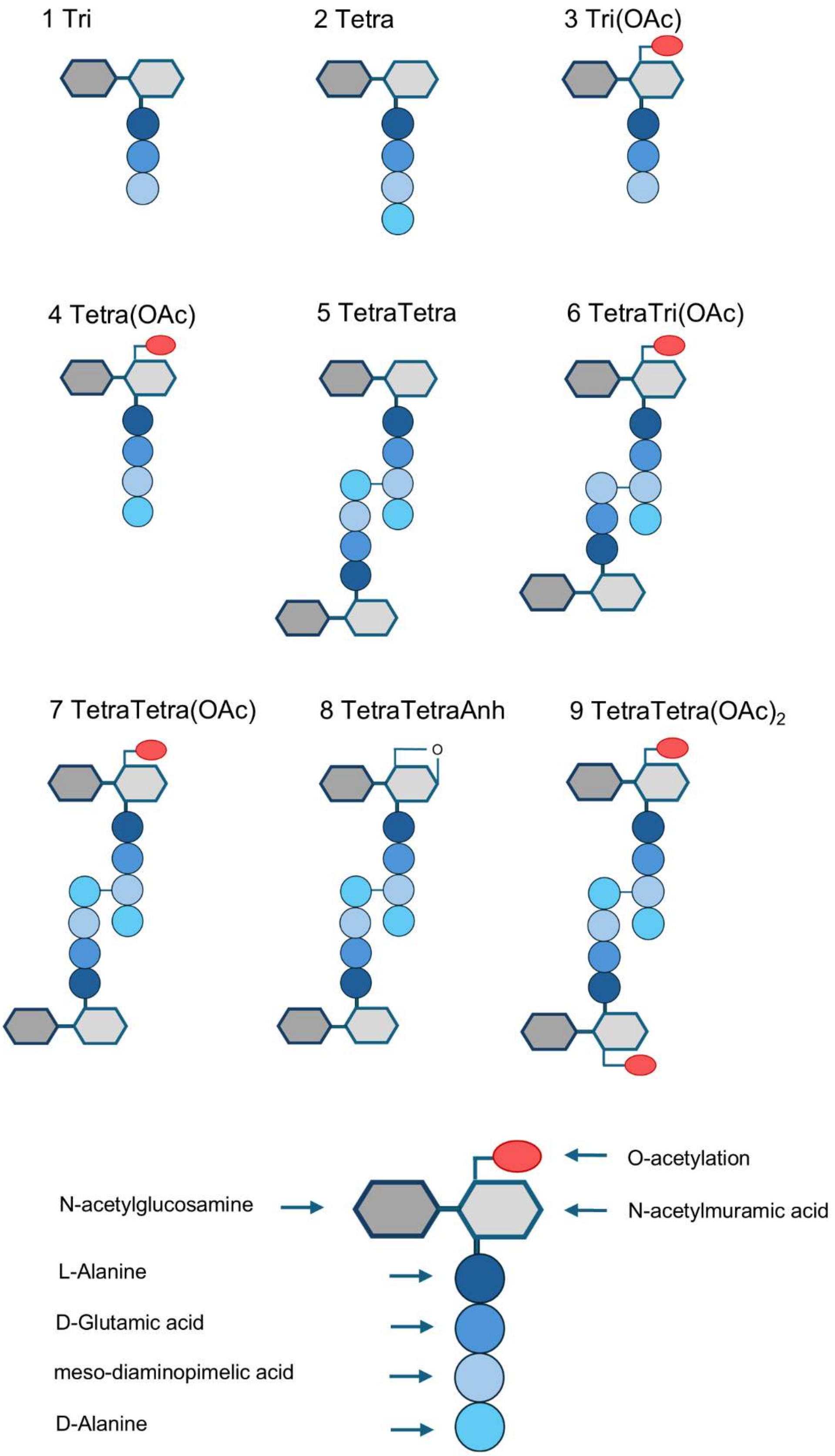


Fig. S4 Schematic structures of muropeptides released from *Z. mobilis* cells. The number corresponds to peak numbers in Fig. 4A, Fig. S3A, and Fig. S6.

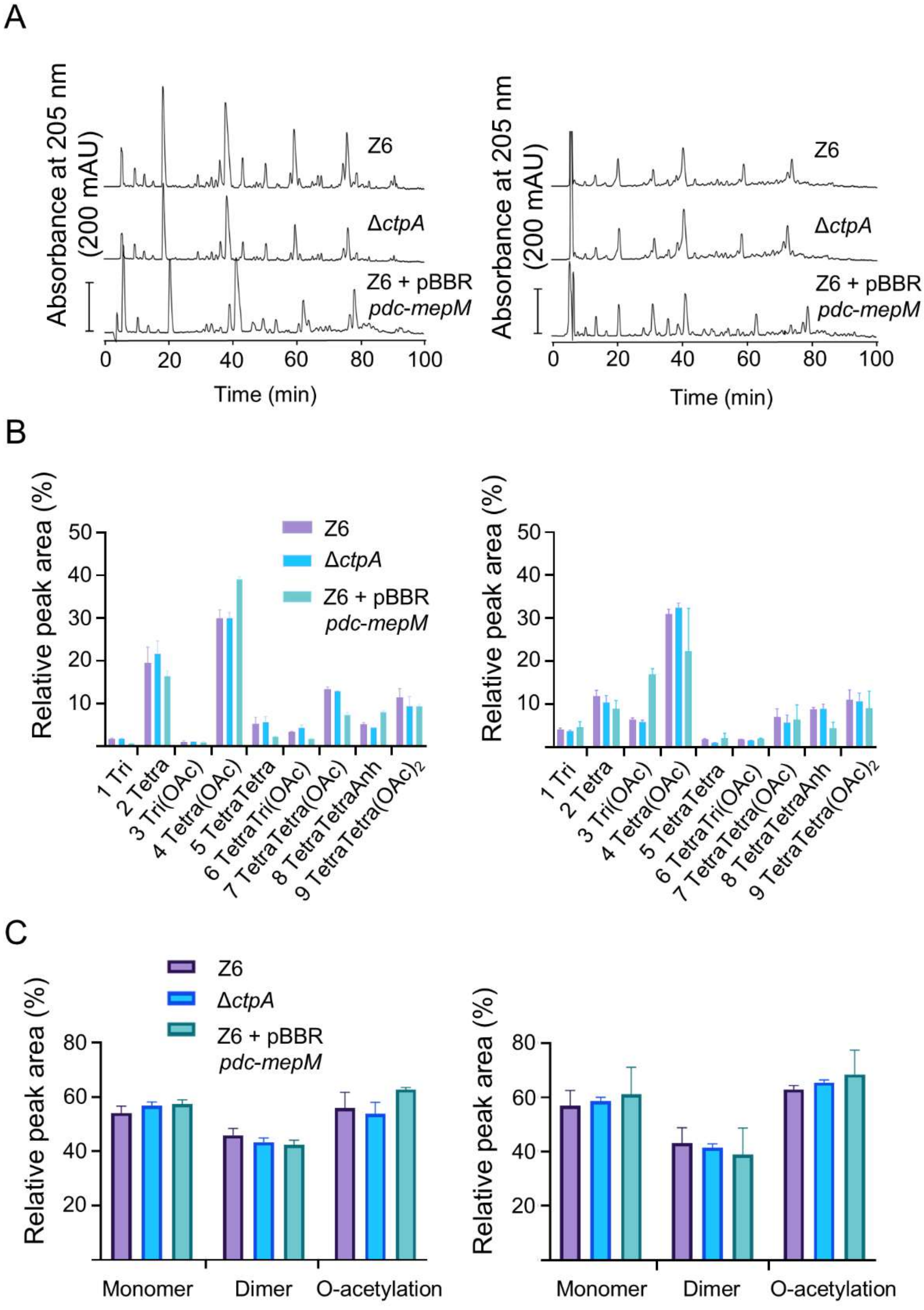


Fig. S5. Muropeptide profile of *Z. mobilis* strain Z6 (wild-type), Δ*ctpA* and Z6 *pdc-mepM*.

(A) Cells of Z6 (wild-type), Δ*ctpA* and Z6 *pdc-mepM* were grown under regular (left) and salt (right) conditions, and their PG were isolated. The muropeptides were released by cellosyl, reduced by sodium borohydride and separated by HPLC under O-acetyl group preserving conditions. The chromatograms of Z6 and Δ*ctpA* are duplicate from Fig 4A. (B) Relative quantification of muropeptides from Z6 and Δ*ctpA* grown under no-salt (left) and salt (right) conditions. (C) Relative proportion of monomeric, dimeric and O-acetylated muropeptides. Colours indicate the strains/growth condition.

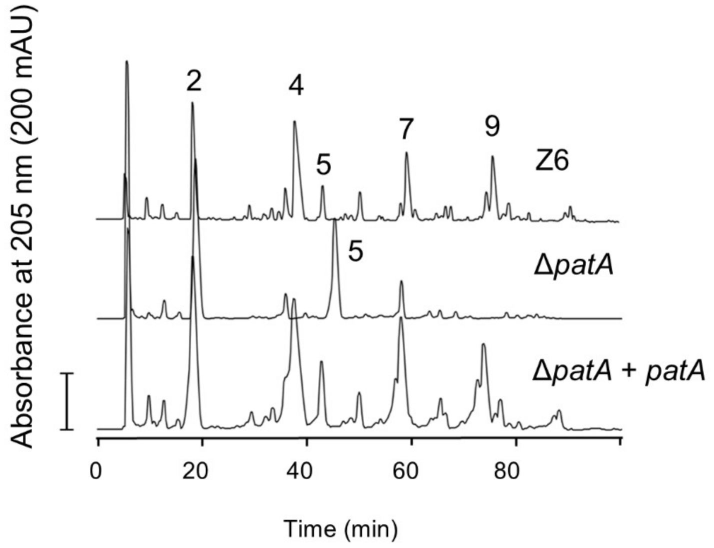


FIg.S6. Muropeptides profile of *Z. mobilis* strain Z6 (wild-type), Δ*patA* and Δ*patA* + *patA* (complemented strain). Cells of three strains were grown under regular growth conditions. The muropeptides from the cells were released by cellosyl, reduced by sodium borohydride and separated by HPLC under O-acetyl group preserving conditions. Be noted that the retention time of Peak 4 in Δ*patA* was shifted as this sample was run separately from the others. Muropeptides are numbered and their structures shown in Fig. S4.

### Supplementary tables

Table S1. Bacterial strains and plasmids used in this study.

| Bacterial strains | Genotype/ description/reference |
|---|---|
| *Zymomonas mobilis* Z6 | ATCC29191 purchased from DSMZ |
| *Escherichia coli* Dh5a | Cloning strain (Lab stock) |
| *Escherichia coli* WM6026 | Conjugation strain (2) |
| *Zymomonas mobilis* Δ*ctpA* | Δ*ctpA* (this study) |
| *Zymomonas mobilis* Δ*ctpA* + *ctpA* | Δ*ctpA* + *ctpA* (this study) |
| *Zymomonas mobilis* Δ*patA* | Δ*patA* (this study) |
| *Zymomonas mobilis* Δ*patA* + *patA* | Δ*patA* + *patA* (this study) |
| *Zymomonas mobilis* pBBR | Z6 + pBBR (this study) |
| *Zymomonas mobilis* pBBR *pdc-mepM* | Z6 + pBBR *pdc-mepM* (this study) |
| *Zymomonas mobilis* pBBR *pdc-mepM* (H287A) | Z6 + pBBR *pdc-mepM* (H287A) (this study) |

Table S2. Oligos used in this study.

| Plasmids | description/reference |
|---|---|
| pKK15534 | Suicide vector (2) |
| pKK15534 + *sacB* | pKK15534 carrying *B. subtilis sacB* (this study) |
| pKK15534 Δ*ctpA* | pKK15534 carrying Δ*ctpA* cassettes (this study) |
| pKK15534 + *ctpA* | pKK15534 + *sacB* carrying *ctpA* insertion cassettes (this study) |
| pKK15534 Δ*patA* | pKK15534 carrying Δ*patA* cassettes (this study) |
| pKK15534 + *patA* | pKK15534 + *sacB* carrying *patA* insertion cassettes (this study) |
| pBBR | Lab stock |
| pBBR *pdc-mepM* | pBBR carrying *pdc* promoter and *mepM* (this study) |
| pBBR *pdc-mepM* (H287A) | pBBR *pdc-mepM* with the mutation H287A (this study) |

Table S3. A list of enhanced proteins in Δ*ctpA* under salt conditions.

| Accession | Gene | gene annotation | log FC2 | P-value |
|---|---|---|---|---|
| AFN56367.1 | ZZ6_0468 | cell division protein FtsL | 1.61207698 | 0.00000005 |
| AFN57534.1 | ZZ6_1677 | endopeptidase MepM | 1.61082639 | 0.00000011 |
| AFN57370.1 | ZZ6_1505 | septum formation initiator DivIC | 1.26758016 | 0.00000279 |
| AFN56779.1 | ZZ6_0886 | hypothetical protein transporter | 1.18367341 | 0.00000608 |
| AFN56793.1 | ZZ6_0900 | ammonium transporter | 1.13329781 | 0.00003357 |
| AFN56550.1 | ZZ6_0653 | flagella basal body P-ring formation protein FlgA | 1.03044958 | 0.00035359 |
| AFN57348.1 | ZZ6_1483 | tonB-dependent siderophore receptor | 0.97573066 | 0.00353524 |
| AFN56826.1 | ZZ6_0933 | PepSY-associated TM helix domain protein | 0.94149198 | 0.00008456 |
| AFN57302.1 | ZZ6_1437 | extensin family protein | 0.90445173 | 0.00000368 |
| AFN57009.1 | ZZ6_1122 | methyl-accepting chemotaxis sensory transducer | 0.90052160 | 0.00033675 |
| AFN56612.1 | ZZ6_0717 | signal transduction histidine kinase | 0.89670457 | 0.00045371 |
| AFN57373.1 | ZZ6_1508 | phosphodiesterase I | 0.87069136 | 0.00006401 |
| AFN56081.1 | ZZ6_0177 | phosphatidate cytidylyltransferase | 0.86008336 | 0.00299904 |
| AFN57484.1 | ZZ6_1625 | hypothetical protein | 0.84415919 | 0.00403771 |
| AFN56594.1 | ZZ6_0699 | succinate dehydrogenase membrane anchor subunit | 0.83931356 | 0.00184612 |
| AFN57548.1 | ZZ6_1691 | gamma-glutamyltransferase | 0.83708535 | 0.00000225 |
| AFN56142.1 | ZZ6_0238 | cellulose synthase catalytic subunit | 0.83625218 | 0.00412790 |
| AFN56312.1 | ZZ6_0411 | protein of unknown function DUF192 | 0.83507995 | 0.00104589 |
| AFN56649.1 | ZZ6_0754 | phosphoesterase PA-phosphatase | 0.81428437 | 0.00074947 |
| AFN56237.1 | ZZ6_0336 | purine nucleoside permease | 0.79774888 | 0.02641677 |
| AFN56944.1 | ZZ6_1056 | hypothetical protein | 0.79732337 | 0.00003339 |
| AFN57397.1 | ZZ6_1535 | TonB-dependent receptor | 0.76353785 | 0.00033841 |
| AFN57255.1 | ZZ6_1390 | hypothetical protein | 0.75923528 | 0.00027323 |
| AFN56030.1 | ZZ6_0125 | potassium transport system protein Kup | 0.75355910 | 0.00000074 |
| AFN57022.1 | ZZ6_1136 | carbohydrate-selective porin OprB | 0.75043935 | 0.00002096 |
| AFN56294.1 | ZZ6_0393 | capsular polysaccharide transport system permease | 0.74682896 | 0.00000069 |
| AFN56660.1 | ZZ6_0765 | GtrA family protein | 0.72507108 | 0.00010170 |
| AFN56238.1 | ZZ6_0337 | Xanthine/uracil/vitamin C permease | 0.72035548 | 0.00015460 |
| AFN56074.1 | ZZ6_0170 | CDP-diacylglycerol/serine O-phosphatidyltransferase | 0.70684025 | 0.00011966 |
| AFN56585.1 | ZZ6_0690 | sodium:dicarboxylate symporter | 0.69601679 | 0.00007877 |
| AFN56327.1 | ZZ6_0427 | hopene-associated glycosyltransferase HpnB | 0.69451356 | 0.03835694 |
| AFN56912.1 | ZZ6_1024 | TonB-dependent siderophore receptor | 0.66952645 | 0.00093194 |
| AFN56965.1 | ZZ6_1077 | S1/P1 nuclease | 0.65941698 | 0.00042556 |
| AFN57019.1 | ZZ6_1132 | major facilitator superfamily MFS_1 | 0.64927937 | 0.01473404 |
| AFN57540.1 | ZZ6_1683 | hypothetical protein | 0.64912631 | 0.00210692 |
| AFN56863.1 | ZZ6_0973 | hypothetical protein | 0.64420395 | 0.00002604 |
| AFN56171.1 | ZZ6_0268 | phosphate ABC transporter | 0.63982345 | 0.00006993 |
| AFN57344.1 | ZZ6_1479 | lytic transglycosylase MltA | 0.63079272 | 0.00148274 |
| AFN56508.1 | ZZ6_0611 | ATP synthase subunit | 0.62623706 | 0.00102737 |
| AFN56329.1 | ZZ6_0429 | cation diffusion facilitator family transporter | 0.61363286 | 0.00000177 |
| AFN55993.1 | ZZ6_0088 | hypothetical protein | 0.61053082 | 0.02115910 |
| AFN56217.1 | ZZ6_0316 | peptidase S10 serine carboxypeptidase | 0.59359605 | 0.00014308 |
